# Telomere damage induces internal loops that generate telomeric circles

**DOI:** 10.1101/2020.01.29.924951

**Authors:** Giulia Mazzucco, Armela Huda, Martina Galli, Daniele Piccini, Michele Giannattasio, Fabio Pessina, Ylli Doksani

**Affiliations:** IFOM, the FIRC Institute of Molecular Oncology, via Adamello 16, 20139 Milan, Italy; Dipartimento di Oncologia ed Emato-Oncologia, Università degli Studi di Milano, Via Festa del Perdono 7, Milan 20122, Italy

## Abstract

Extrachromosomal telomeric circles are commonly invoked as important players in telomere maintenance, but their origin has remained elusive. Using electron microscopy analysis on purified telomeres we show that, apart from known structures, telomeric repeats accumulate internal loops (i-loops), that occur in proximity of nicks and single-stranded DNA gaps. I-loops are induced by single-stranded damage at normal telomeres and represent the majority of telomeric structures detected in ALT (Alternative Lengthening of Telomeres) tumor cells. Our data indicate that i-loops form as a consequence of the exposure of single-stranded DNA at telomeric repeats. Finally, we show that these damage-induced i-loops can be excised to generate extrachromosomal telomeric circles resulting in loss of telomeric repeats. Our results identify damage-induced i-loops as a new intermediate in telomere metabolism and reveal a simple mechanism that links telomere damage with the accumulation of extrachromosomal telomeric circles and telomere erosion.

## Introduction

Mammalian telomeres are made of several kilobases of tandem TTAGGG repeats that are required to protect chromosome ends from the DNA damage response. Erosion of telomeric repeats can lead to senescence and genome instability and therefore plays important roles in ageing and tumorigenesis ^1^. Extrachromosomal circular DNA made of telomeric repeats (t-circles) have been found in a wide range of organisms and are thought to play opposite roles in telomere maintenance. They have been associated to telomere loss via deletion/trimming of telomeric repeats, while in some contexts, like in ALT cells they could promote telomere elongation through rolling-circle amplification^2–4^. C-circles (t-circles with a covalently closed and partially single-stranded C-rich strand) accumulate in ALT cells and provide a diagnostic marker for ALT tumors^5^. Despite their relevance in telomere biology, it is not clear how these circles are generated. Telomeric circles have been detected in cells expressing a TRF2 mutant that lacks the N-terminal basic domain and, given the role of TRF2 in the formation/maintenance of t-loops, it has been proposed that t-circles could form via nucleolytic excision of the t-loop structure^2,6,7^. However, more recently t-circles have been found also in normal cells and in an ever-growing list of mutants, apparently unrelated to t-loop metabolism, or to each other, suggesting the existence of alternative mechanisms for their formation^3,8–14^for a review see^15^. T-circles can be detected through a rolling-circle replication assay, and their presence is often inferred from the appearance in two-dimensional agarose gel electrophoresis (2D-gels) of an arc, compatible with the migration of relaxed circular DNA^2,8,16^. Telomeric circles have been found in electron microscopy (EM) imaging of ALT telomeres^17^but further analysis, (e.g. identification of intermediates of t-circle formation) has been hampered by the inconsistency of available procedures for telomere purification.

Using a newly-developed telomere purification procedure, combined with EM analysis, we found that damaged telomeres tend to form internal loops (i-loops), likely due to the exposure of single-stranded DNA at telomeric repeats. These structures migrate in the t-circle arc in 2D-gels and represent the majority of telomeric structures found in ALT cells. We show that damage-induced i-loops can be excised as telomeric circles, resulting in telomere loss. These results identify damage-induced i-loops as a key intermediate in telomere circle formation and provide a mechanism that links telomere damage with t-circle formation and telomere erosion.

## Results

### A two-step procedure for the purification of mammalian telomeres

Telomeric repeats lack restriction sites and this property has been exploited for their enrichment by digestion of non-telomeric DNA with frequent cutters and then purification of large DNA fragments, containing telomeres, in gel filtration columns^18,19^. Key telomere features (e.g. t-loops, t-circles) have been visualized with this approach, however, incomplete digestion of non-telomeric DNA and poor fractionation of milligrams of DNA in gel filtration columns, have limited its applications. To overcome these issues, we developed a two-step procedure for the large-scale purification of telomeric repeats from mammalian cells. First, 2.5 mg of genomic DNA are digested with frequent cutters and separated in a sucrose gradient (Figure 1A). Then, high molecular weight fractions, containing the telomeric repeats are collected, digested again with a new mixture of restriction enzymes (see materials and methods) and separated in a preparative agarose gel (Figure 1B). The high molecular weight DNA recovered from the agarose gel shows ~1000-fold increase in telomeric repeats signal compared to the starting material, while more abundant mouse long interspersed repeats (BamHI repeats) are undetectable (Figure 1C). Telomere enrichment was confirmed in single-molecule IF-FISH analysis, where over 80% of the DNA molecules from enriched samples are recognized by a telomeric probe, while less than 1 in 1000 telomeric fibers were present in non-enriched samples (Figure 1D). We obtained similar enrichment levels from human cells with long telomeres (Figure S1A, B).

**Figure 1.**
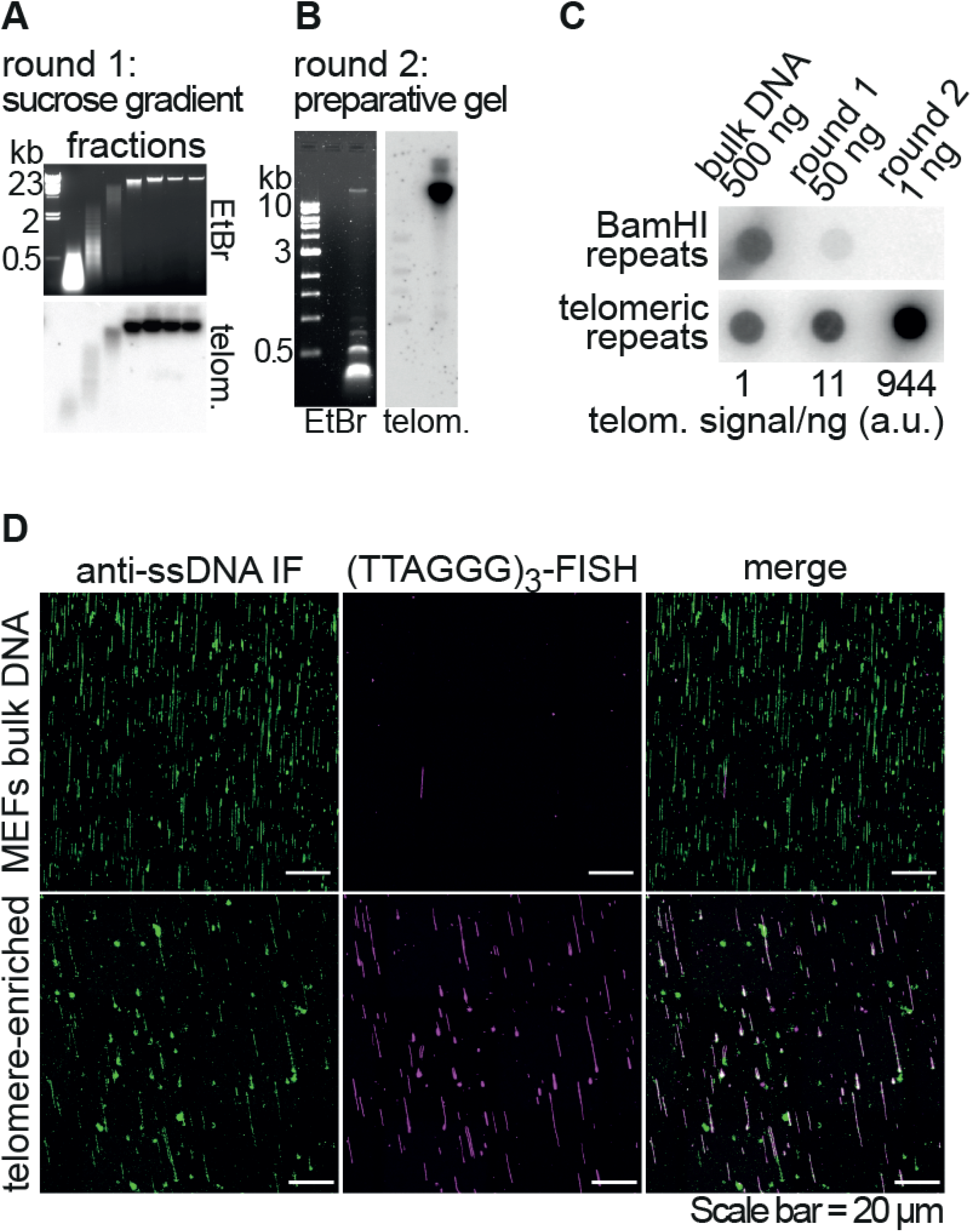
A two-step procedure for the purification of mammalian telomeres. **A.** Top: agarose gel showing the separation of the large telomeric repeat fragments from the bulk DNA in a sucrose gradient. Genomic DNA (~2.5 mg) from SV40-MEFs was digested with HinfI and MspI. The digested DNA was separated by centrifugation on a sucrose gradient. Seven fractions were collected and an aliquot (~1/500) of each fraction was loaded on an agarose gel. Bottom: the gel was blotted onto a membrane and hybridized with a TTAGGG repeats probe to verify that telomeric repeats remained in the high molecular weight (HMW) fractions. **B.** Left: agarose gel showing the separation of the large telomeric repeat fragments from the remaining non-telomeric DNA, in the second purification round. The HMW DNA, contained in the last four fractions of the sucrose gradient described in (A), was recovered and digested with RsaI, AluI, MboI, HinfI, MspI, HphI and MnlI. The digested DNA was separated on a preparative agarose gel and the DNA migrating in the area above 5 kb was extracted from the gel. The image shows an aliquot (~1/100) of the digested DNA, separated on an agarose gel. Right: the gel was blotted onto a membrane and hybridized with a TTAGGG repeats probe to verify that telomeric repeats remained in the HMW area. **C.** Dot blot analysis showing the enrichment of telomeric repeats. The indicated amounts of DNA from each enrichment step were spotted on a membrane and hybridized either with a probe recognizing the long interspersed BamHI repeats or TTAGGG repeats. The amount of TTAGGG repeat signal/ng was quantified and reported relative to the signal/ng value in the initial, non-enriched DNA. **D.** Single molecule analysis showing the enrichment of the telomeric repeats. The DNA was combed onto silanized coverslips, denatured in situ and labeled sequentially with an antibody against single-stranded DNA and a Cy3-labeled (TTAGGG)_3_PNA probe.

### Frequent i-loops in telomere-enriched samples

We employed the procedure described above to isolate telomeres from mouse embryo fibroblasts (MEFs) and analyze their structure in EM. The DNA was crosslinked with psoralen in vivo, prior to cell lysis, and the telomere-enriched material was spread with the BAC method and rotary shadowed with Platinum^20^. In telomeric spreads, DNA fragments ranged from 2 to 40 kb. As expected, telomeric samples were enriched in t-loops, although their absolute frequency in our spreads was lower than in previous settings (Figure S2A-C)^19,21^.

One salient feature we observed in telomeric spreads was the occurrence of molecules with i-loops (Figure 2A). Differently from t-loops, which sequester one end of the DNA molecule and are therefore terminal, i-loops appeared as crossings of the internal regions of the molecules, where the ends are not engaged. In three independent experiments, with SV40-immortalized MEFs, around 14% of molecules in the telomere-enriched samples had one or more i-loops. In control spreads of genomic DNA, fragmented at a similar size by restriction digestion, i-loops occurred in around 3% of the molecules (Figure 2B). An abundance of molecules with i-loops was seen also in telomeric spreads from HeLa 1.3 cells with long telomeres (Figure S3A-C).

**Figure 2.**
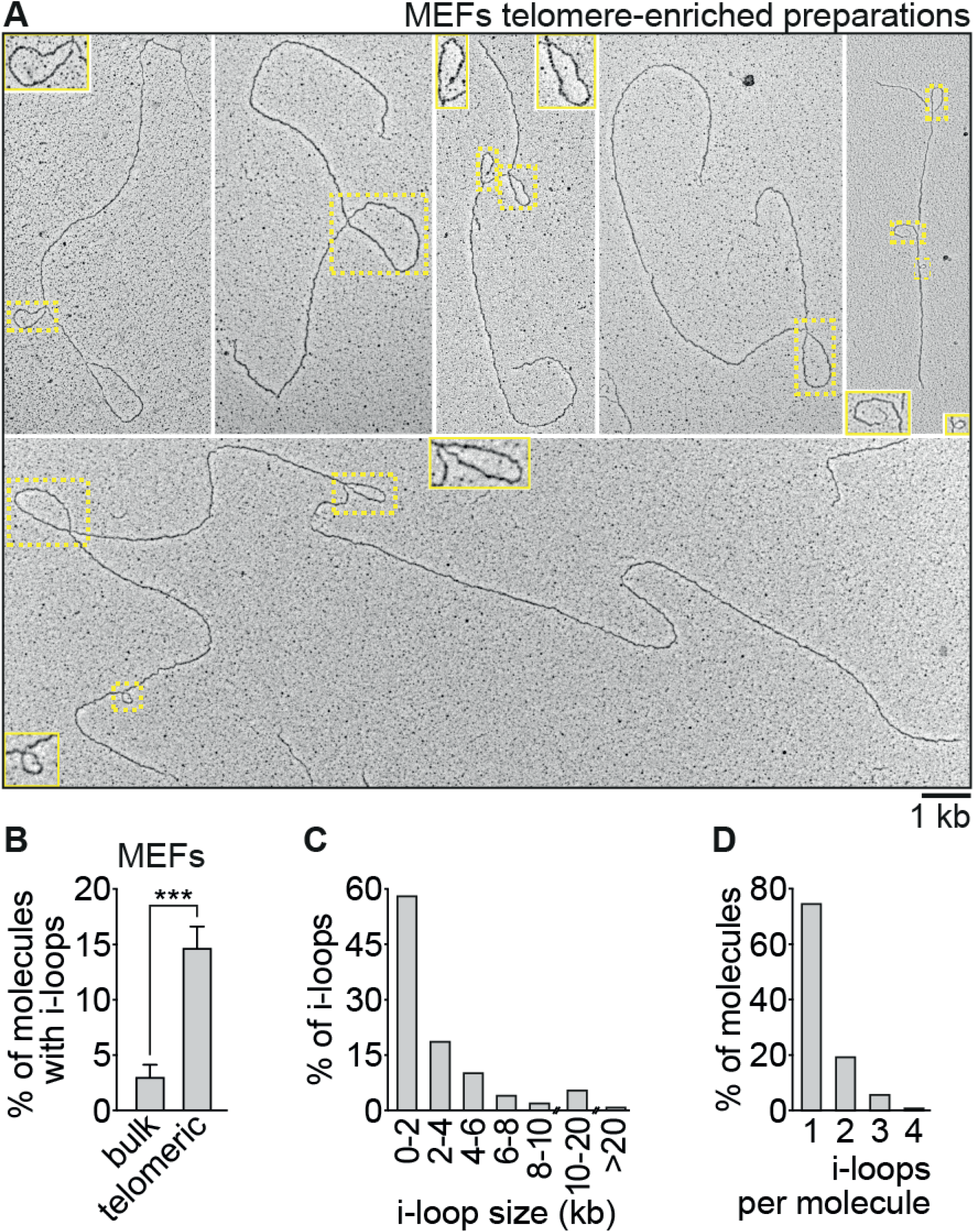
Frequent i-loops in telomere-enriched samples. **A.** Examples of molecules with i-loops found in the telomeric samples. Telomeric DNA, enriched with the procedure described in Figure 1, was analyzed in transmission Electron Microscopy (EM). I-loops are indicated by the yellow rectangles. Insets show 2X enlargements of some of the areas inside the yellow rectangles. **B.** Quantification of i-loop occurrence. Telomere-enriched (telomeric) and non-enriched (bulk) DNA was analyzed in EM as described above. The non-enriched, bulk DNA was digested with KpnI, which generates fragments of 10 kb in average. The fraction of molecules containing one or more i-loops was scored in 3 independent experiments (n=310, 715, and 845 molecules for telomere-enriched samples and 1466, 776 and 1776 molecules for non-enriched samples). Error bars represent standard deviation. P value was derived from unpaired, two-tailed, Student’s t-test. **C.** Size distribution of the i-loops in the telomeric DNA. N = 342 i-loops. **D.** Distribution of the number of i-loops per molecule in the telomeric DNA. N = 266 molecules with i-loops.

I-loops ranged from 0.2 to 25 kb, with a median size of 1.6 kb (Figure 2C). In the majority of cases i-loops occurred once per molecule, but in about 25% of cases two or more i-loops were present on the same molecule (Figure 2D). We examined, at higher magnification, the structure of the i-loops and noticed that about one in four had a thinner, apparently single-stranded, region at the junction (Figure 3). In another 20% of the loops, a short gap and/or a small flap was visible in one of the DNA strands near the junction. Based on these observations, we hypothesized that i-loops could represent structural transitions that occur at sites of single-strand damage (i.e. nicks and gaps) on the telomeric repeats.

**Figure 3.**
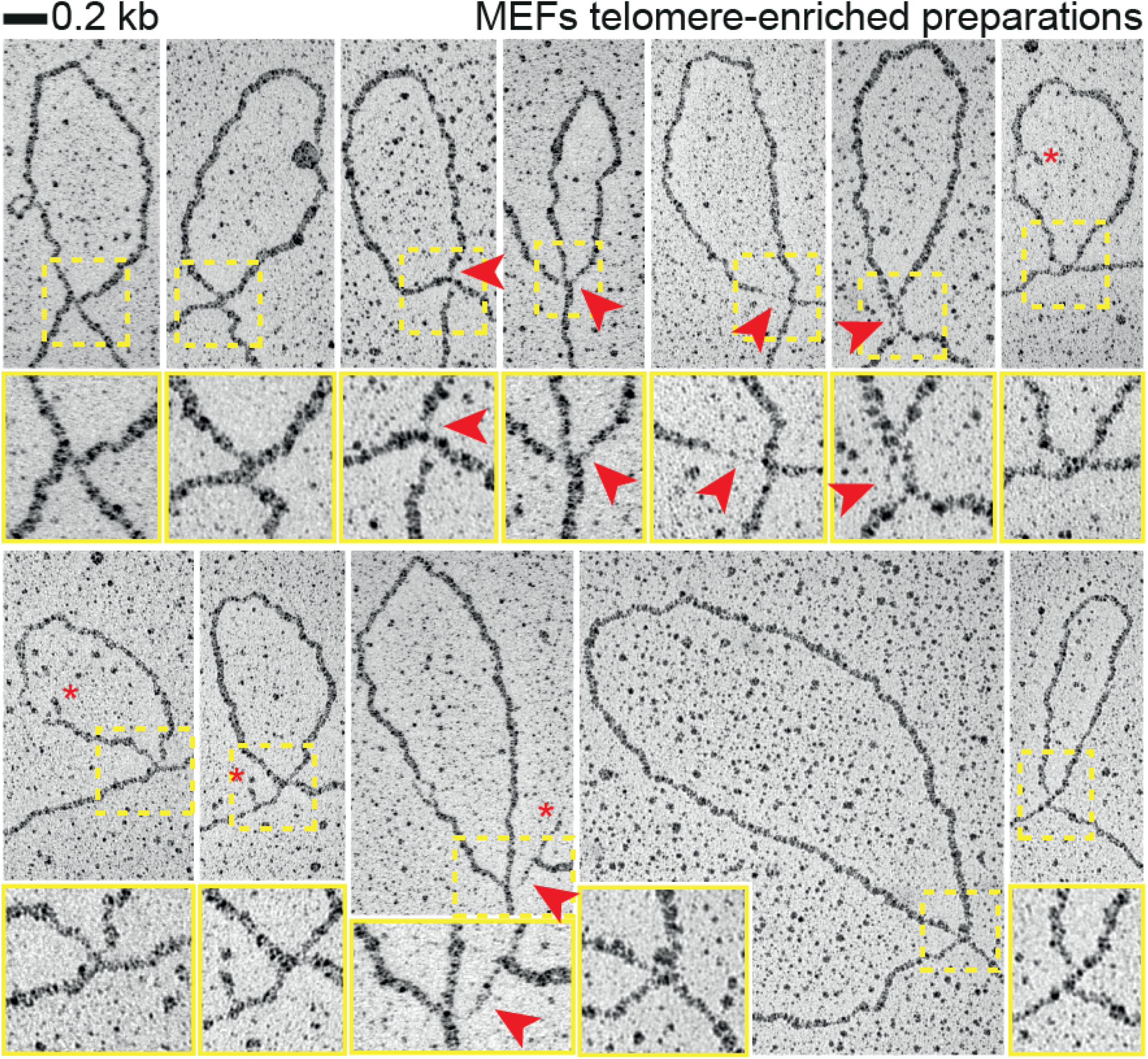
I-loops often occur in proximity of strand damage. High magnification images of i-loops observed in telomere-enriched fractions of the experiment described in Figure 2. A 2X enlargement of the area inside the yellow rectangle is shown under each loop. Red arrows indicate regions of single-stranded DNA at the loop junction, while red asterisks indicate small flaps in proximity of the loop junction.

### I-loops are the majority of telomeric structures detected at ALT telomeres

Since i-loops often occurred in proximity of single strand damage, we turned our attention to ALT cells, which contain nicks and gaps at telomeres and show unusual telomeric structures in 2D-gels^2,22^. In particular, a faint, slow-migrating arc is detected in 2D-gels at ALT telomeres; this signal is commonly known as the t-circle arc and is attributed to the presence of extrachromosomal telomeric circles^2,16,22^. Although the t-circle arc is compatible with the migration of relaxed circular DNA, there is no direct evidence on the types of telomeric structures that populate it. We decided to purify the DNA molecules from the t-circle area of the 2D gel in order to visualize their structure in EM. Genomic DNA was prepared from U2OS cells and telomeres were enriched with the same procedure described above, except that in the second round of enrichment the DNA was separated in 2D-gel (Figure 4A, Figure S4A). The areas of the second-dimension gel containing the t-circle arc and the linear telomeres were excised (Figure 4A), the DNA was recovered and analyzed in EM. As expected, the material purified from the t-circle area was richer in DNA structures, although it still contained substantial amounts of linear fragments (Figure S4B). Around 9% of the molecules were circular and in 28% of these circles single-stranded gaps were visible (Figure 4B, C). This result confirms previous reports on the presence of double-stranded and partially single-stranded telomeric circles in ALT cells^2,5,17^. However, molecules with one or more i-loops represented 40% of all DNA recovered from the t-circle area, over 4-fold more abundant than telomeric circles (Figure 4B, D, Figure S4B). Therefore, i-loops represent the vast majority of telomeric structures identified by 2D-gels in ALT cells. Also at ALT telomeres, i-loops often occurred in proximity of strand damage (Figure 4D).

**Figure 4.**
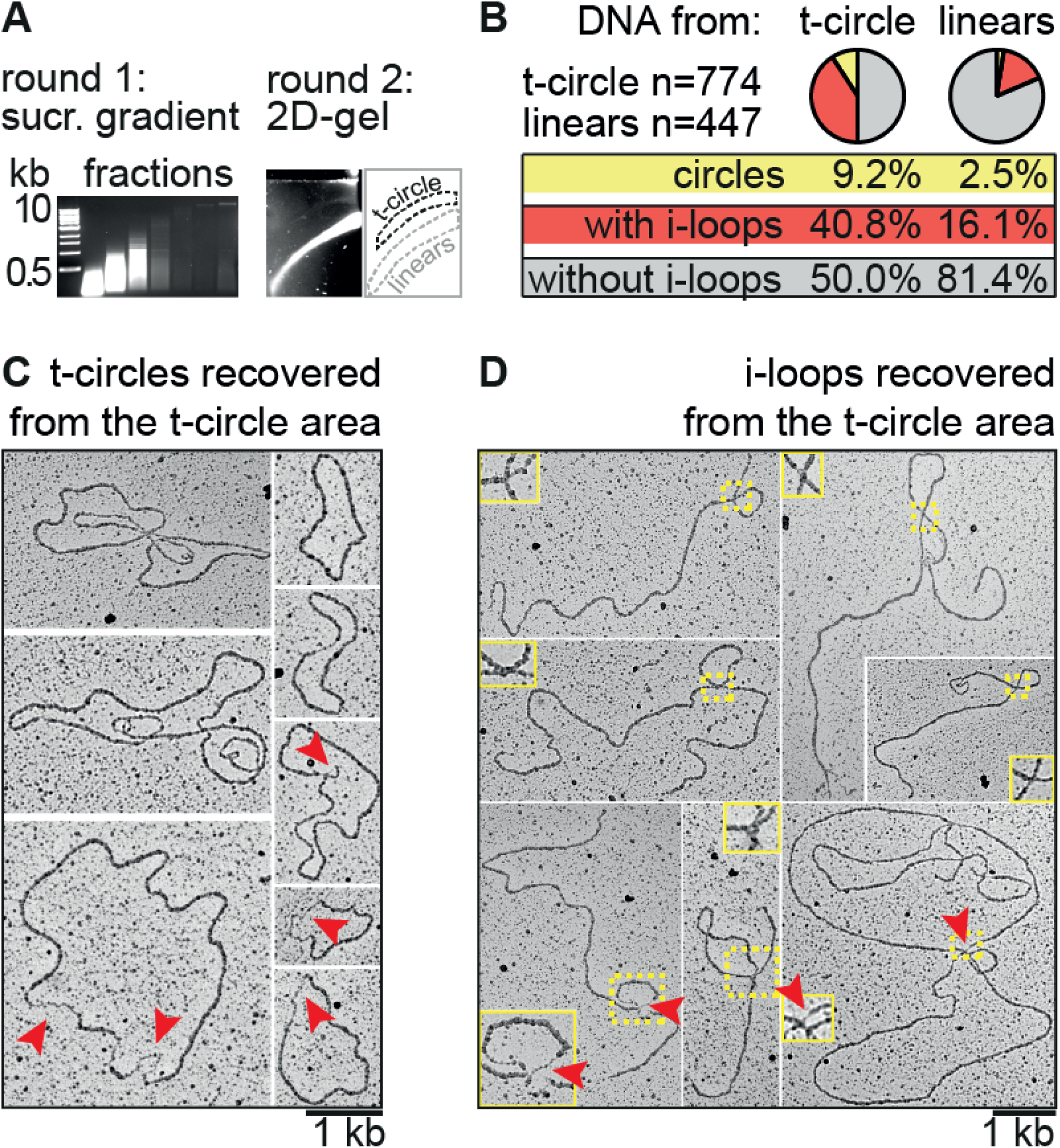
I-loops are the majority of telomeric structures detected at ALT telomeres. **A.** Procedure for the purification of telomeric DNA migrating in the linear and the t-circle arc of ALT telomeres. Genomic DNA (~2.5 mg) extracted from U2OS cells was processed for the telomere enrichment procedure as described in Figure 1A. The HMW DNA, contained in the last 4 fractions of the sucrose gradient, was collected, digested again as described in in Figure 1B and separated in a 2D-gel. A strong linear signal and a faint t-circle arc were visible in the second-dimension gel (right). These areas, were excised and the DNA was recovered from the gel. **B.** Pie chart showing the distribution of the molecules recovered from the 2D-gel. Percentages of the 3 major categories are shown. Note that i-loops can occur also in molecules having a t-loop at the end, or at branched molecules. This sub-distribution is reported in Figure S4B. **C.** Example of circular molecules found in the DNA purified from the t-circle arc. Arrows indicate regions of single-stranded DNA. **D.** Examples of i-loops found in the DNA purified from the t-circle arc. Insets represent 2X enlargements of the areas inside the yellow rectangles. Red arrows indicate regions of single-stranded DNA at the loop junction.

### I-loops are induced by single-strand damage at telomeric repeats

Since i-loops were associated with single-stranded telomere damage and populated the t-circle arc in U2OS cells, we asked whether telomere damage alone can induce their formation and therefore the appearance of the t-circle arc in 2D-gels. To test this hypothesis, we used mild DNase I treatment, that introduces both nicks and short single-stranded gaps on DNA^23^. MEFs nuclei were incubated with increasing concentrations of DNase I, then genomic DNA was isolated, digested with frequent cutters and separated in 2D-gels. Telomeres from mock-treated nuclei migrated mainly as linears with no t-circle arc visible, while damaged telomeres, isolated from nuclei that were treated with DNase I, showed a strong accumulation of the t-circle arc (Figure 5A). We obtained the same result in human cell lines with long or short telomeres, although t-circle arc induction strongly decreased with telomere length (Figure S5A). No arc was induced at the abundant mouse BamHI repeats, or in the bulk genomic DNA, showing that, at these magnitudes, this is not a general feature of nicked DNA (Figure S5B).

**Figure 5.**
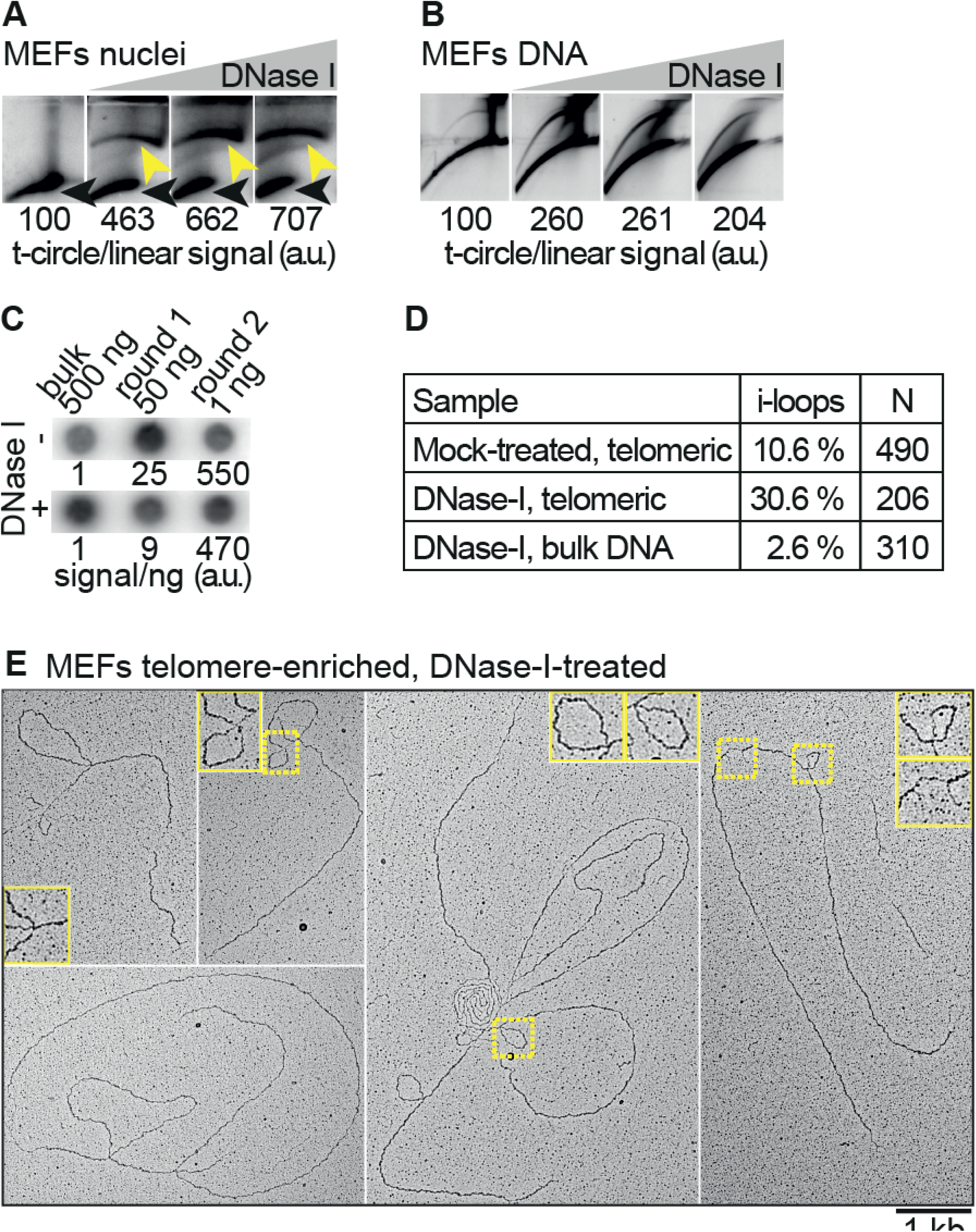
I-loops are induced by single-strand damage at telomeric repeats (see also Figure S5). **A.** 2D-gel analysis showing that the t-circle arc can be strongly induced by formation of nicks and gaps at telomeres. MEFs nuclei were incubated with either 0; 1; 2.5 or 5 μg/ml of DNase I for 8 minutes at RT. The reaction was stopped and the genomic DNA was isolated. 5 μg were digested with AluI and MboI and separated on 2D-gels. The gels were blotted on a membrane and hybridized with a TTAGGG repeats probe. The ratio of the telomeric signal in the t-circle arc (yellow arrows) and in the linears (black arrows), is reported relative to the untreated sample, which was arbitrarily set to 100. **B.** 2D-gel analysis showing that the t-circle arc can form spontaneously, in the presence of nicks and gaps at the telomeric repeats. Isolated mouse genomic DNA was incubated with either 0; 0.1; 0.2 or 0.4 μg/ml of DNase I for 8 min at RT. The reaction was stopped, the DNA was extracted with phenol-chloroform, digested with AluI and MboI, separated on 2D-gels, blotted on a membrane and hybridized with a probe recognizing the TTAGGG repeats. The ratio of the telomeric signal in the t-circle arc and in the linears, is reported relative to the untreated sample which was arbitrarily set to 100. **C.** Dot blot showing the enrichment of the telomeric repeats, after the large-scale DNase I treatment. Around 500 × 10^6^SV40-MEFs nuclei were incubated either with 0 or 5 μg/ml of DNase I for 8 minutes at RT. The reaction was stopped, genomic DNA was isolated and telomeres were enriched with the procedure described in Figure 1. The indicated amounts from each enrichment step were spotted on a membrane and hybridized with a probe recognizing the TTAGGG repeats. The telomeric signal per ng of DNA is reported relative to the non-enriched DNA. **D.** Accumulation of i-loops at telomeres damaged by DNase I. Telomere-enriched DNA from the experiment described in (C) was analyzed in EM. The percentage of molecules with internal loops is reported. A KpnI-digested bulk DNA control was included for the sample treated with DNase I. **E.** Examples of molecules with internal loops observed at telomere preparations, from DNase I-treated nuclei. Insets show 2X enlargements of the area inside the yellow rectangles.

Since the DNase I treatment was performed on isolated nuclei, we asked if the chromatin environment or any chromatin-associated factor is required for the generation of the structures migrating in the t-circle arc. Surprisingly, mild DNase I treatment of isolated, protein-free, genomic DNA resulted in a strong induction of the t-circle arc at telomeric repeats, while the same structural transition was not observed in the bulk DNA, or at the BamHI repeats (Figure 5B, Figure S5C). The absence of specialized enzymatic activities in this setting rules out the accumulation of telomeric circles (see also below), demonstrating that the 2D-gel arc, commonly-identified as the t-circle signal, can be generated in the absence of telomeric circles.

Having established that telomere strand damage is sufficient for the induction of the t-circle arc, we verified that this process was indeed associated with the accumulation of telomeric i-loops, as predicted. The same DNase I treatment described above, was performed in large scale, then genomic DNA was isolated, processed through the two-step procedure for the enrichment of telomeric repeats and analyzed in EM (Figure 5C). Telomere strand damage induced by DNase I resulted in a 3-fold increase in i-loops, while in the bulk DNA control i-loops remained at basal levels (Figure 5D, E). Together these experiments show that, accumulation of nicks and gaps at telomeric repeats is sufficient to induce i-loops and generate the t-circle signal in 2D-gels.

Based on these experiments, we hypothesized that i-loops are formed via spontaneous annealing and branch migration events, occurring at sites of gaps or nicks favored by the extremely high abundance of homology at telomeric repeats. If this is the case, then formation of i-loops after the induction of DNA damage should be limited by inter-strand psoralen crosslinking that prevents DNA branch migration and preserves native DNA structures^24^. To test this hypothesis, we first performed psoralen crosslinking on the nuclei, before the DNase I treatment and then analyzed telomere structures in 2D-gels, as above. DNase I damaged to a similar extent both crosslinked and non-crosslinked DNA, however, formation of i-loops was strongly inhibited by psoralen crosslinking, as seen by the reduced intensity of the t-circle arc in 2D-gels (Figure 6A). This result suggests that i-loop formation, after the induction of telomere strand damage requires DNA branch migration. Importantly, once formed, i-loops are not sensitive to crosslinking. Indeed, the t-circle arc of U2OS cells was not affected by psoralen crosslinking (Figure 6B), showing that ALT telomeres, which experience endogenous damage, contain i-loops in vivo, prior to psoralen crosslinking.

**Figure 6.**
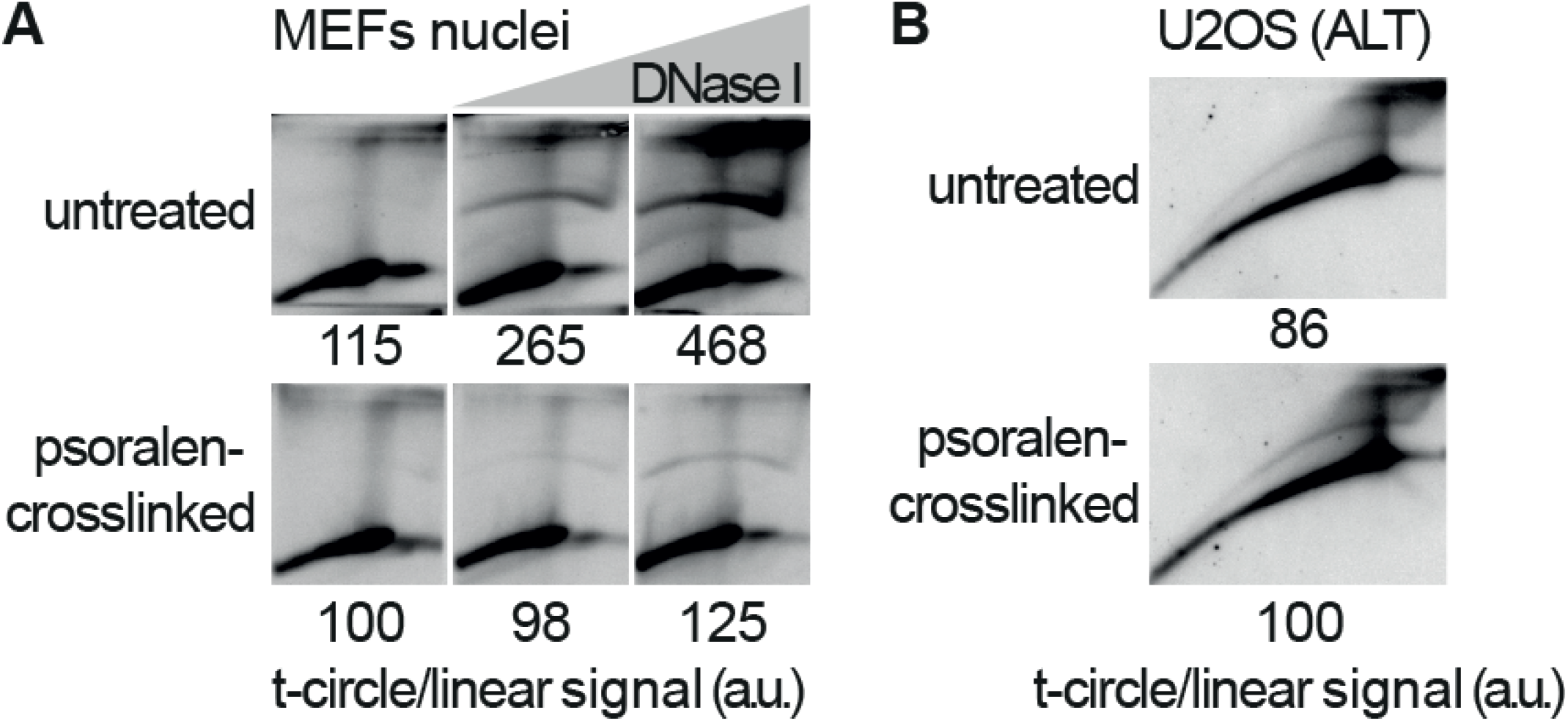
I-loop formation following the induction of DNA damage requires branch migration. **A.** A preparation of SV40-MEFs nuclei was split in two and one half was psoralen crosslinked on ice (in this prep, DNA branch migration is largely prevented). Then, both preps were treated with DNase I and processed for 2D-gels as described in Figure 5A. The ratio of the telomeric signal in the t-circle arc and in the linears, is reported relative to the untreated and crosslinked sample, which was arbitrarily set to 100. **B.** 2D-gel analysis showing that the intramolecular loops that accumulate in ALT cells are not affected by psoralen crosslinking. U2OS cells were psoralen crosslinked on ice, then genomic DNA was extracted and processed for 2D-gels as above. The ratio of the telomeric signal in the t-circle arc and in the linears, is reported relative to the crosslinked sample which was arbitrarily set to 100.

### I-loops are a substrate for the generation of extrachromosomal telomeric circles

The short (TTAGGG) telomere repeat motif, provides a context where almost any single stranded gap exposes intramolecular homology. For instance, two gaps on opposite telomeric strands could generate an i-loop simply by strand annealing (Figure 7A). Further branch migration and strand exchange events could generate a double Holliday Junction (HJ) at the base of the i-loop. Similarly, two gaps on the same telomeric strand could undergo strand exchange and generate i-loops with a single HJ at the base (Figure 7B). We hypothesized that these damage-induced i-loops would be a substrate for HJ resolvases, in a reaction similar to the one proposed for unprotected t-loops^2,6^. In this resolution reaction 50% of events would result in telomere deletions and generation of extrachromosomal telomeric circles (Figure 7C). In order to test whether i-loops are biochemically a substrate for the generation of telomeric circles, we first treated genomic DNA with DNase I, to induce i-loops at telomeres, then incubated the DNA with a nuclear extract from HeLa cells (Figure 7D). The DNA was then recovered and subjected to the rolling-circle replication assay^5^. Neither DNase I treatment nor incubation with the nuclear extract alone induced telomeric circles, while the combination of the two resulted in a 5-fold increase in circles (Figure 7D, F). Importantly accumulation of telomeric circles required both Mg++ and ATP, consistent with the requirements of a Holliday junction resolution reaction. This experiment shows that i-loops are a substrate for the generation of extrachromosomal telomeric circles, in a reaction resembling HJ resolution.

**Figure 7.**
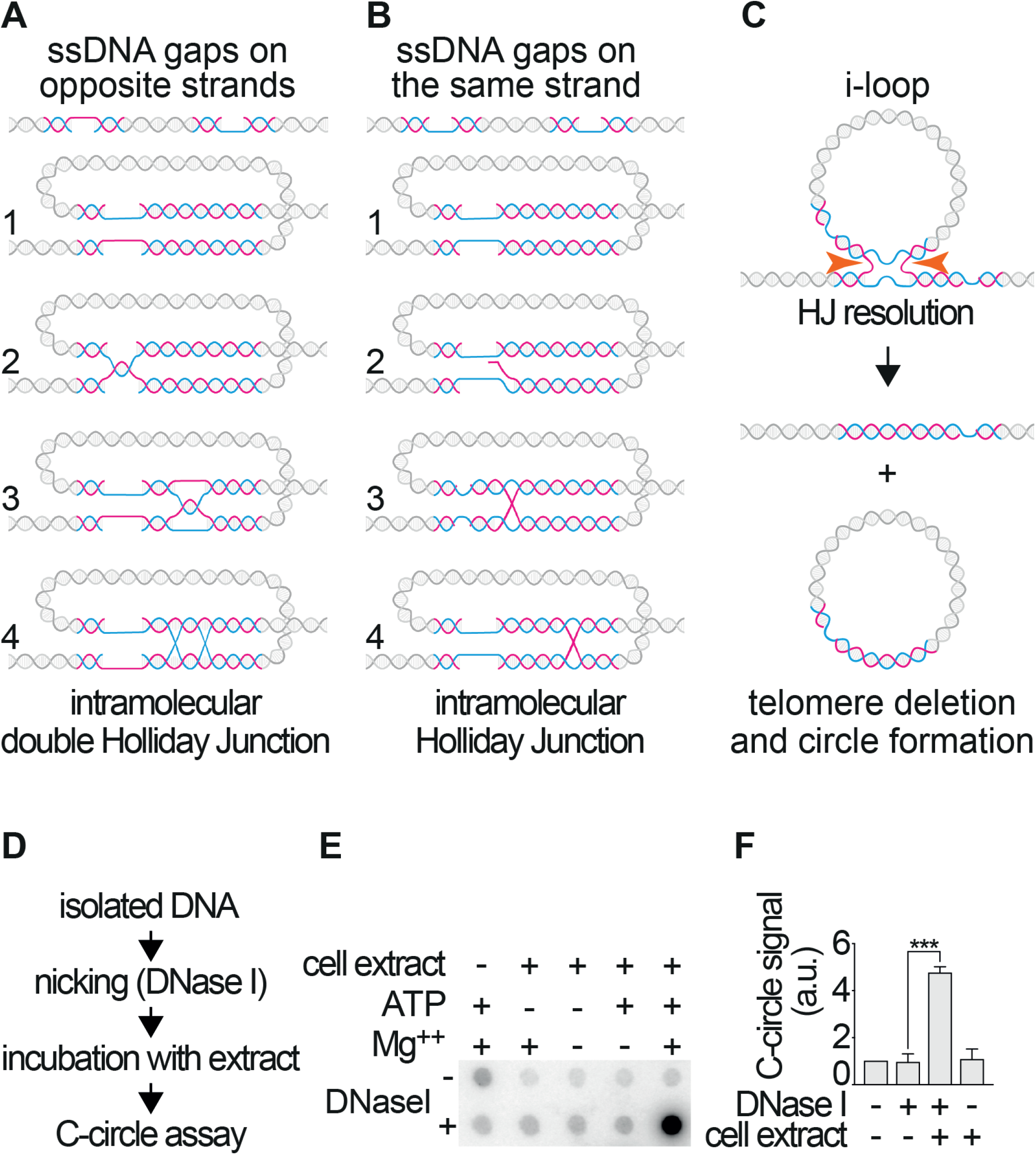
I-loops are a substrate for the generation of extrachromosomal telomeric circles. **A.** Model for the formation of i-loops, in the presence of short single-strand gaps on opposite telomeric strands (i.e. one gap on the G-strand and one on the C-strand of the same molecule). Exposed complementary DNA can come in close proximity by looping and undergo base pairing (step 1). Plectonemic pairing can occur simply by strand rotation, resulting in the formation of an i-loop that will resemble a DNA knot (step 2). The loop junction could branch migrate as a hemicatenane (step 3) that could be transformed in a double Holliday junction by the pairing of the opposite strands (step 4). **B.** Model for the formation of i-loops, in the presence of short single-strand gaps on the same telomeric strand (i.e. both gaps on the G or on the C strand of the same molecule). The gaps can come in close proximity by DNA looping (step 1) and promote an exchange of the complementary strands (step 2) resulting in an intramolecular loop with a single Holliday junction at the base (step 3), that can undergo branch migration (step 4). **C.** Model for the generation of the telomeric circles via the excision of i-loops. An i-loop with a Holliday Junction at the base becomes a substrate for Holliday Junction resolvases. Cleavage on the horizontal axis of the image (orange arrows) will result in the excision of the loop as a circle and telomere loss. Note that the excised circle, would contain a nick, resulting from the HJ resolution and a single-stranded gap (one of the original gaps that induced formation of the i-loop). **D.** Schematic representation of the experimental procedure used to test the model shown in (C). Isolated DNA was nicked with low concentrations of DNase I, which induces the formation of i-loops at telomeres. The reaction was stopped, DNA was extracted and incubated for 30 minutes at 37°C with a HeLa nuclear extract in the presence of Mg++ and ATP, to allow HJ resolution. The DNA was then purified and the presence of telomeric circles was assayed with the C-circle assay. **E.** Dot blot analysis of the circle assay of the experiment described in (D). The DNA from the C-circle assay was blotted on a membrane and hybridized with a probe recognizing the TTAGGG repeats. A strong C-circle signal accumulates only in the combined treatment nicking and incubation with the extract. Quantification of the C-circle signal from 3 independent experiments as the one described in (D). The signal is reported relative to the untreated sample (no DNase I, no extract) which was set to 1. P value was derived from unpaired, two-tailed, Student’s t-test.

## Discussion

We identify damage-induced i-loops as key intermediates that link telomere damage with telomere erosion and the generation of extrachromosomal telomeric circles. These results predict that conditions associated with chronic telomere (or DNA) damage (e.g. chemotherapeutics; replication stress), will favor the formation of telomeric circles and telomere loss, while factors that prevent formation of i-loops at sites of damage (e.g. factors that prevent strand exchange or improper single strand annealing at telomeres) would counteract the accumulation of extrachromosomal telomeric circles. Given that ALT cells are known to experience endogenous telomere damage, the mechanism proposed in Figure 7 could help explain the continuous generation of telomeric circles in ALT cells. In the same view, accumulation of telomeric damage, could be a common denominator that explains the presence of t-circles in many mutants in genes involved in DNA metabolism and telomere maintenance^9–13^. Frequent formation of i-loops could provide yet another challenge to replication fork progression at telomeres and contribute to telomere fragility^25^. I-loops could be a relevant substrate for specialized helicases like RTEL1, Blm and WRN, which could prevent formation or promote branch migration/dissolution of i-loops at telomeric repeats, thereby reducing the probability of telomere loss due to i-loop excision^26–28^.

Damage-induced i-loops might occur also at other tandem repeats, explaining the formation of circular DNA at these sequences from yeast to human^16,29,30^. In this view, it is important to notice that the overall rate of i-loop formation will be higher at repetitive elements with a shorter repeated motif, because they will be more likely to expose complementary sequences when damaged. Therefore, telomeres, with a repeat unit of 6 nt, will be more prone to generate extrachromosomal circles compared to most other long repeats. This high propensity of telomeric repeats to form i-loops that can be excised as circles, would result in continuous and stochastic variations in the number of repeats thus explaining, at least in part, the amplitude of telomere length heterogeneity across different chromosomes or different cells.

A positive correlation between telomere length and accumulation of the t-circle signal in 2D-gels has been reported in normal and stem cells, indicating the existence of a trimming mechanism that controls telomere length^3,31^. Our results suggest that, as telomere length increases so will the probability of i-loop formation and excision due to stochastic damage. This correlation could be relevant in understanding the sources of dysfunctional telomeres and how telomere length evolves in different organisms.

## Acknowledgements

We are grateful to Titia de Lange, for allowing YD to initiate this project in her lab, for reagents, and helpful discussions. We thank Marco Foiani and Fabrizio d’Adda di Fagagna for technical support and helpful discussions and Elia Zanella for helpful discussions. We are grateful to members of the IFOM Cell Biology Unit, for their invaluable assistance with growing large cell cultures. We thank Paolo Maiuri for developing an ImageJ macro for the annotation and storage of the EM pictures and the IFOM imaging facility for technical assistance. YD lab is supported by the Associazione Italiana per la Ricerca sul Cancro, AIRC, IG 19901. FP is supported by the Fondazione Telethon GGP17111.

## Author contributions

AH and DP assisted with the setting of the sucrose gradient conditions for the telomere enrichment procedure. AH performed the U2OS telomere purification from 2D-gels. M.Giannattasio provided technical assistance with the initial DNA spreading and EM procedure. M.Galli performed the EM experiment in HeLa 1.3 cells. FP assisted with the HeLa extract incubation. GM and YD performed the rest of the experiments. YD conceived the study and wrote the paper.

## Methods

### Cell culture

SV40LT-immortalized MEFs were grown in D-MEM (Lonza, BE12-614F) supplemented with 10% fetal bovine serum (EuroClone, ECS0180L), 2 mM L-glutamine (EuroClone, LOBE17605F), 100 U/ml penicillin-0.1 μg/ml streptomycin (EuroClone, ECB3001L), 0.1 mM non-essential amino acids (Microtech, X-0557). HeLa 1.3 cells were grown in D-MEM (Lonza, BE12-614F) supplemented with 10% fetal bovine serum (EuroClone, ECS0180L), 2 mM L-glutamine (EuroClone, LOBE17605F), 100 U/ml penicillin-0.1 μg/ml streptomycin (EuroClone, ECB3001L).

### Enrichment of telomeric repeats

Around 500×10^6^cells, were harvested and resuspended in ice-cold PBS. For psoralen crosslinking, the cell suspension was poured in a 10 cm dish and kept on ice while stirring, throughout the procedure. The suspension was first incubated with 30 μg/ml 4, 5’, 8-trimethylpsoralen (Sigma, T6137, stock 2 mg/ml in DMSO, stored at −20°C) for 5 minutes in the dark and then exposed to 365 nm UV light for 8 minutes in a UV Stratalinker 1800, (Stratagene), with 365 nm UV bulbs (model UVL-56, UVP) at 2-3 cm from the light source. The incubation and irradiation steps were repeated three more times (4 cycles total). Cells were then lysed in TNES buffer (Tris 10 mM pH8.0, NaCl 100 mM, EDTA 10 mM; 0.5% SDS) incubated with 50 μg/ml RNaseA (Sigma, R500) for 60 min at 37°C, and then with 100 μg/ml Proteinase K (Roche, 3115887001) for 12 hours at 37°C. The DNA was extracted with Phenol Chloroform Isoamyl alcohol 25:24:1 (Sigma, P2069) followed by an extraction with Choloroform (VWR, 22711) and precipitation with isopropanol. Around 2.5 mg of DNA was digested overnight with 750 units of HinfI and MspI (NEB). The digestion was precipitated and loaded on a sucrose gradient, 10%-20%-30% sucrose, 8 ml each fraction, in TNE buffer and centrifuged in SW32-Ti rotor (Beckman) at 30100 rpm (111265 g) for 16 hours. The HMW fractions containing the telomeric repeats were collected, concentrated and washed twice with Tris 10mM pH 8.0 in Amicon Ultra-15 Ultracel-PL PLTK, 30 kDa MWCO (Millipore/MERCK UFC903024) filters. The DNA was then digested overnight with 50 units each of RsaI, AluI, MboI, HinfI, MspI, HphI, MnlI (NEB) and then separated on a 0.7% low-melting agarose gel (SeaPlaque Agarose, Lonza, 50100), without ethidium bromide. Fragments migrating above the 5 kb band of the marker were extracted using the Silica Bead DNA gel extraction kit (Thermo Fisher Scientific, K0513) following the manufacturer’s instructions, except that once the DNA was bound, the beads were not resuspended to avoid mechanical shearing of the DNA. The DNA was eluted in TE 1X and quantified using Qubit dsDNA HS assay kit (Invitrogen, Q32854).

### Single molecule analysis of telomeric enrichment

Around 10 ng of bulk genomic DNA or telomere-enriched DNA was combed on silanized coverslips (Genomic Vision, COV-002) using the DNA Fiber Comb apparatus (Genomic Vision, version 3 REF: MSC-001). The coverslips with the DNA were subjected to the following treatments: baking for 2-3 hours at 60°C, denaturing in 0.5 M NaOH, 1M NaCl for 8 min, followed by two washes in PBS and dehydration in ethanol series (70%; 90%; 100%, 1 min each). Blocking with 5% BSA in PBS for 1 hour at 37°C and incubation with an anti single-stranded DNA antibody (Sigma, MAB3034) diluted 1:80 in 5% BSA in PBS, for 2 hours at RT, followed by three washes with PBS −0.05% Tween 20. Incubation with Alexa488-labeled anti-mouse secondary antibody (Invitrogen, A1101) diluted 1:400 in 5% BSA in PBS, followed by three washes with PBS −0.05% Tween 20 and dehydration in ethanol series (70%; 90%; 100%, 1 min each). Incubation with a Cy3-labeled TTAGGG_3_ PNA probe (PNA Bio, F1006), 50 nM in 70% formamide (Thermo Scientific, 17899) 0.5% Blocking reagent (Roche 11096176001) and 10 mM Tris, pH 7.4, for 3 min at 80°C and then 2 hours at RT, followed by 2 x15 min washes in formamide 70%, 10 mM Tris, pH 7.4 and three washes with PBS-0.05% Tween 20. The coverslips were mounted using ProLong Gold (Invitrogen, P36930). Large overlapping areas were acquired and stitched in a DeltaVison microscope.

### Electron microscopy analysis

EM analysis was performed as described in ^20^. Typically, 5 μl of telomere-enriched DNA corresponding to 5-20 ng were used for each spread. For non-enriched controls, 30 ng of KpnI-digested genomic DNA was spread using the same method. The DNA recovered from the linear and t-circle arc of 2D-gels, was spread using the droplet method as in ^32^. Briefly, 1 ng of DNA in 28 μl of TE 1X, was mixed with 30 μl of Formamide (Thermo Scientific, 17899) and 2 μl of benzyldimethyl-alkylammonium chloride (BAC) 0.08%. The droplet was incubated for 5 min at RT and the surface was gently touched with a carbon-coated EM grid, previously activated by contact with an ethidium bromide solution 33 μg/ml in TE1X. The grids were then processed for staining with Uranyl Acetate and rotary shadowing as described in ^20^. TEM pictures were taken using a FEI Tecnai12 Bio twin microscope operated at 120 KV and equipped with a side-mounted GATAN Orius SC-1000 camera controlled by the Digital Micrograph software. Images in DM3 format were analyzed using the ImageJ software. In these conditions 0.36 μm correspond to 1 kb of double-stranded DNA.

### DNase I treatment on isolated nuclei

MEF nuclei were isolated as described previously ^33^. Briefly, cells were collected by trypsinization, washed with ice-cold PBS, and resuspended in ice-cold fibroblast lysis buffer (12.5 mM Tris pH 7.4, 5 mM KCl, 0.1 mM spermine, 0.25 mM spermidine, 175 mM sucrose, supplemented with protease inhibitor cocktail (Roche, 11836170001) at a concentration of 8 × 10^6^cells/ml). After 10 min of incubation on ice, 0.02 vol 10% NP-40 was added and cells were incubated for 5 min on ice. Nuclei were collected by centrifugation at 1000g for 5 min at 4°C and washed once with ice-cold Nuclei Wash Buffer (NWB) (10 mM Tris-HCl pH 7.4, 15 mM NaCl, 60 mM KCl, 5 mM MgCl_2_, 300 mM sucrose) and resuspended in NWB. When indicated, psoralen crosslinking was performed on the nuclei suspension in NWB, as described above for cell suspensions. For the DNase I treatment, 1 volume of nuclei suspension was mixed with 1 volume of DNase I cocktail (NWB supplemented with CaCl_2_ 2 mM, BSA 100 μg/ml, and twice the indicated concentration of DNase I (Roche 10104159001) and incubated for 8 minutes at RT. The reactions were stopped with 0.5 vol of ice-cold stop buffer (50 mM EDTA, 10 mM EGTA). The nuclei were then processed for genomic DNA extraction as described above for cells.

### DNase I treatment on isolated DNA

Genomic DNA, extracted as described above, was incubated with DNase I (Roche 10104159001) in 10 mM Tris-HCl pH 7.4, 15 mM NaCl, 60 mM KCl, 5 mM MgCl_2_, 10 mM CaCl_2_, for 8 min at RT. The reaction was stopped by adding 0.2 vol of EDTA-EGTA 0.25 M each, extracted with 1 volume of phenol-cholorform isoamylalcohol and precipitated in isopropanol.

### 2D-gels

10 μg of genomic DNA was digested overnight with 20 units of AluI and MboI (NEB) and then precipitated with isopropanol. For the analysis of the mouse BamHI repeats, the DNA was digested either with BglI or with KpnI as indicated. The first dimension was run in 0.35% agarose (US-biological, A1015) in TBE 0.5X, without ethidium bromide for 12-24 hours at 1 V/cm. The gel was stained with 0.3 μg/ml ethidium bromide in TBE 0.5X and lanes were excised above 5 kb for mouse, U2OS and HeLa 1.3 telomeres and above 2 kb for HeLa 204 and HTC75 telomeres. The second dimension was run in 0.7% agarose in TBE 0.5X with 0.3 μg/ml ethidium bromide at 3-5 Volts/cm at 4°C. When necessary, psoralen crosslinking was reversed before southern blotting by exposing the gel to 254 nm UV for 10 min in a stratalinker (UVP CL1000 Ultraviolet crosslinker). For Southern blotting, the gel was first incubated 2 × 30 min with the depurination solution (HCl 0.25N) 2 × 30 min with denaturing solution (NaOH 0.5M, NaCl 1.5 M), 2 × 30 min with neutralizing solution (Tris 0.5M pH 7.5, NaCl 3M). The DNA was then transferred by capillarity in SSC 20X onto an Amersham Hybond-X membrane (GE healthcare RPN203). For TTAGGG repeats probe the 800 bp EcoRI fragment of the Sty11 plasmid (a gift from Titia de Lange) ^34^was used. For the BamHI repeats probe a 1 kb EcoRV fragment, containing mouse BamHI dispersed repeats ^35^cloned in pBlue was used. Radioactive signal was captured on phosphor screens (FUJIFILM Storage Phosphor screen MS3543 E), read on a Typhon Trio (GE) and analyzed on ImageJ.

### Incubation with HeLa extracts

1 μg of genomic DNA was incubated with 60 μg of HeLa nuclear extract (6 mg/ml) (IPRACELL, CC012010) in 50mM Tris HCl pH8, 150 mM NaCl, 5 mM MgCl2, 2 mM ATP, 1mM DTT for 35 minutes at 37°C in 20 μl final volume. The reaction was stopped with 0.1 vol of EDTA-EGTA 0.25 M each, extracted with 1 volume of phenol-cholorform isoamylalcohol and precipitated in isopropanol.

### C-circle assay

Was performed according to ^5^. Briefly, 25 ng of genomic DNA, digested with AluI and MboI, were incubated for 12 hours at 30°C with 7.5 Units of Phi29 polymerase (NEB M0269) in Phi29 NEB buffer 1X, supplemented with dNTPs 0,37 mM each, in a final volume of 20 μl. The enzyme was inactivated by heating to 65°C for 20 minutes and the reaction was blotted onto a Hybond-X membrane. Telomeric repeats were detected using the TTAGGG repeats probe described above.

## Supplemental Information

**Figure S1.**
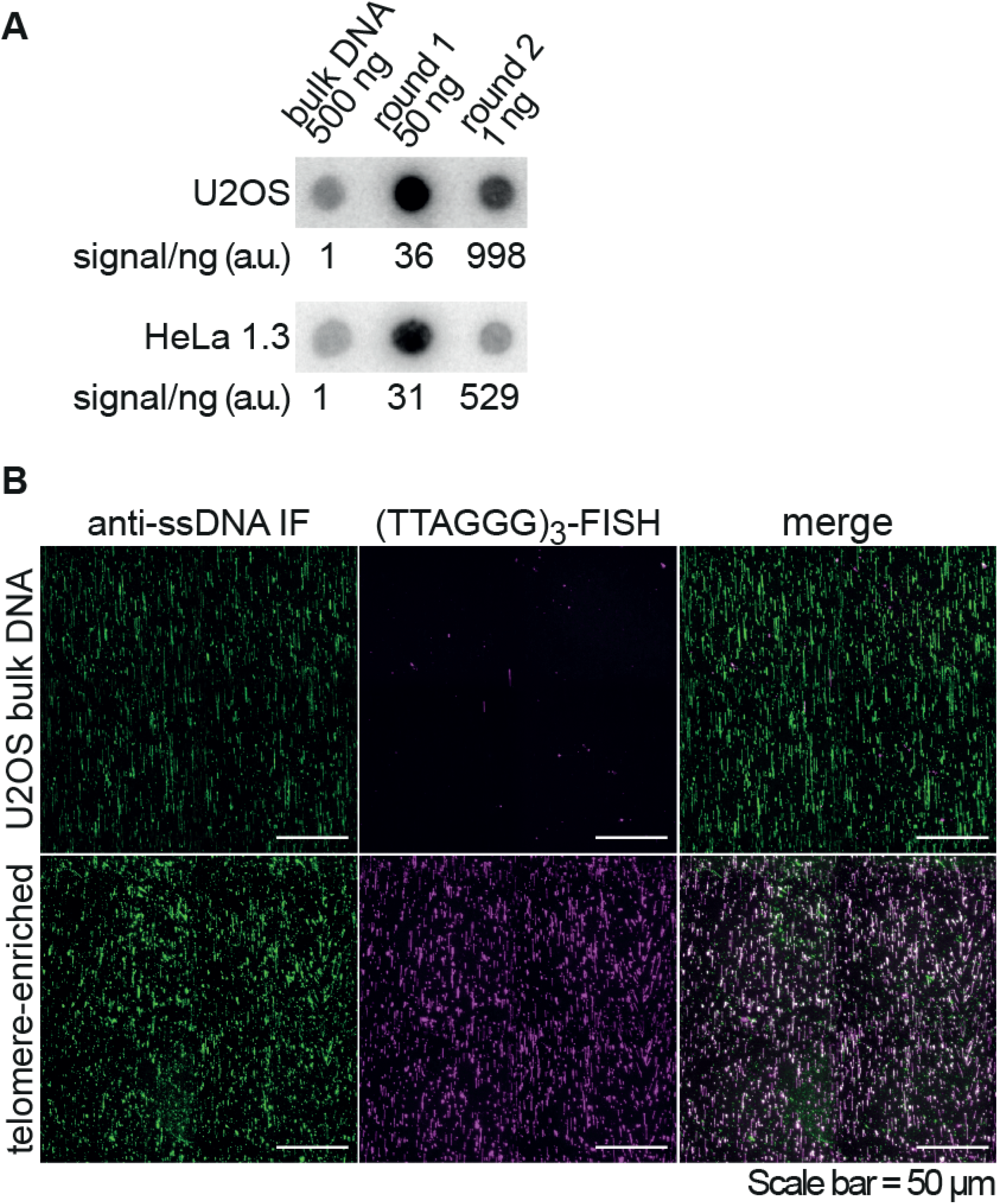
Enrichment of telomeric repeats from human cells (related to Figure 1). **A.** Dot blot analysis showing the enrichment of the telomeric repeats. Genomic DNA prepared from U2OS cells and HeLa 1.3 cells with long telomeres, was subjected to the telomere enrichment procedure (described in Figure 1). The indicated amounts from each enrichment step were spotted on a membrane and hybridized with a probe recognizing the TTAGGG repeats. The amount of signal/ng is reported relative to the non-enriched DNA. **B.** Single molecule analysis showing telomere enrichment from U2OS cells. Enriched telomeric DNA was combed on silanized coverslips, denatured in situ and labeled sequentially with an antibody against single-stranded DNA and a Cy3-labeled (TTAGGG)3 PNA probe.

**Figure S2.**
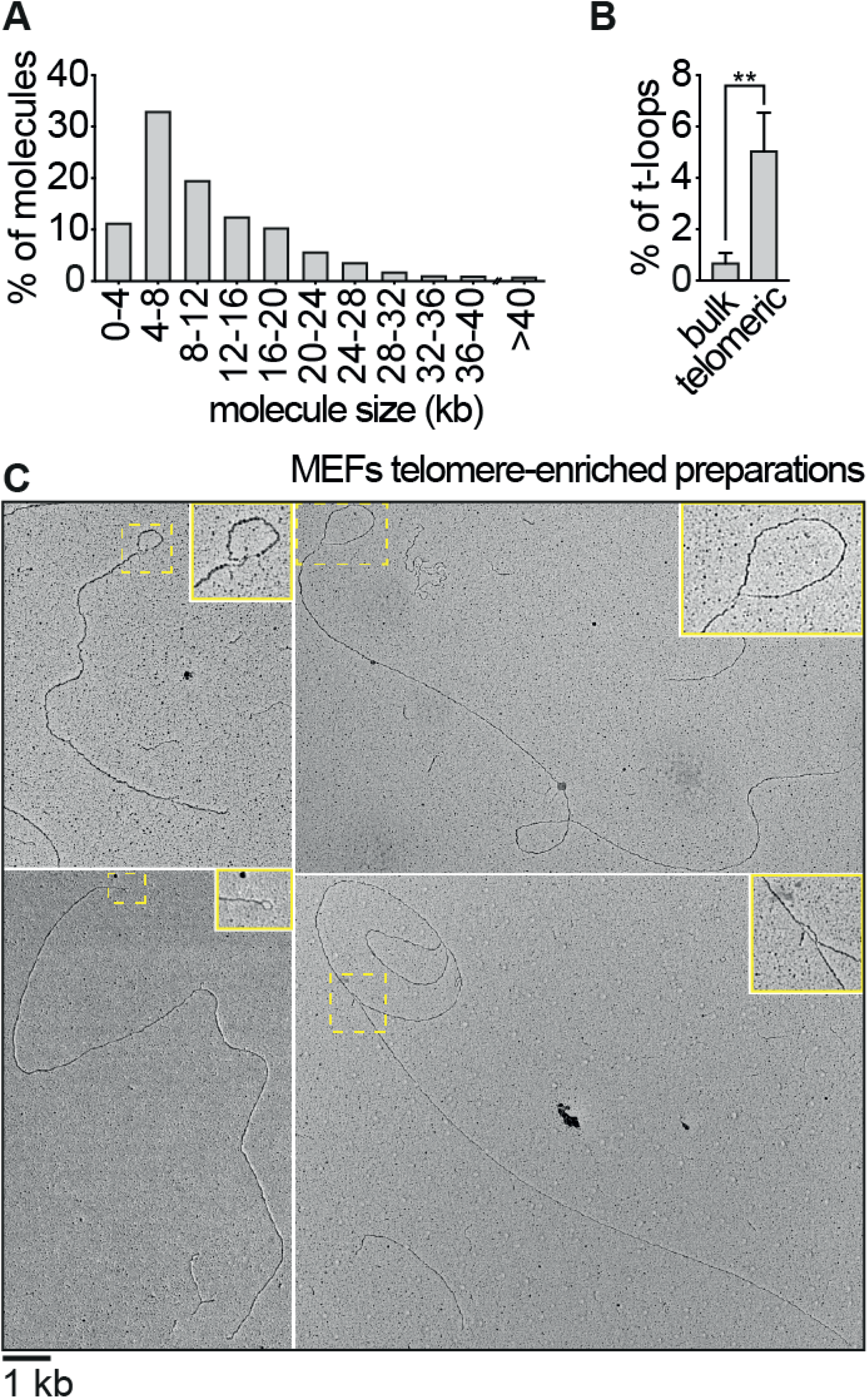
Occurrence of t-loops in the telomere-enriched samples (related to Figure 2). **A.** Molecule length distribution from the experiments with telomere-enriched DNA, described in Figure 2. N=1516 molecules. **B.** Frequency of t-loops in the telomere-enriched (telomeric) and non-enriched (bulk) samples from the experiments described in Figure 2. Error bars represent standard deviation from 3 independent experiments. P value was derived from unpaired, two-tailed, Student’s t-test. **C.** Examples of molecules with t-loops observed in the telomere-enriched samples. Insets show 2X enlargements of the area inside the yellow rectangles.

**Figure S3.**
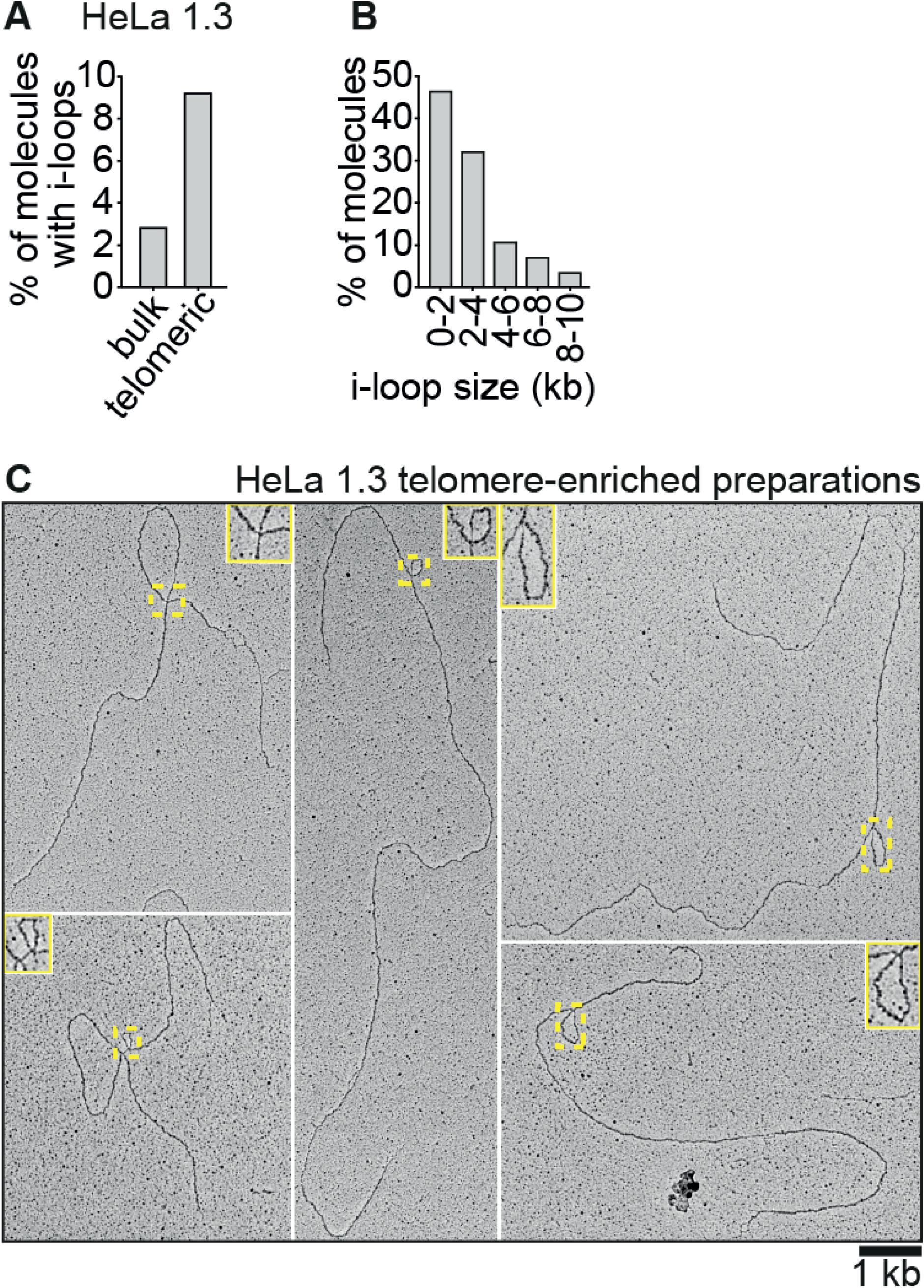
Accumulation of i-loops at human telomeres. **A.** Quantification of intramolecular loop occurrence in telomere-enriched (telomeric) and non-enriched (bulk) DNA from HeLa 1.3 cells with long telomeres. The percentage of molecules with i-loops is reported for each sample. Telomere enriched N=239, non-enriched N=1510 molecules. **B.** I-loop size distribution, from the experiment described in (A). N=28 internal loops. **C**. Examples of molecules with i-loops observed in the telomere-enriched sample from HeLa 1.3 cells. Insets show 2X enlargements of the area inside the yellow rectangles.

**Figure S4.**
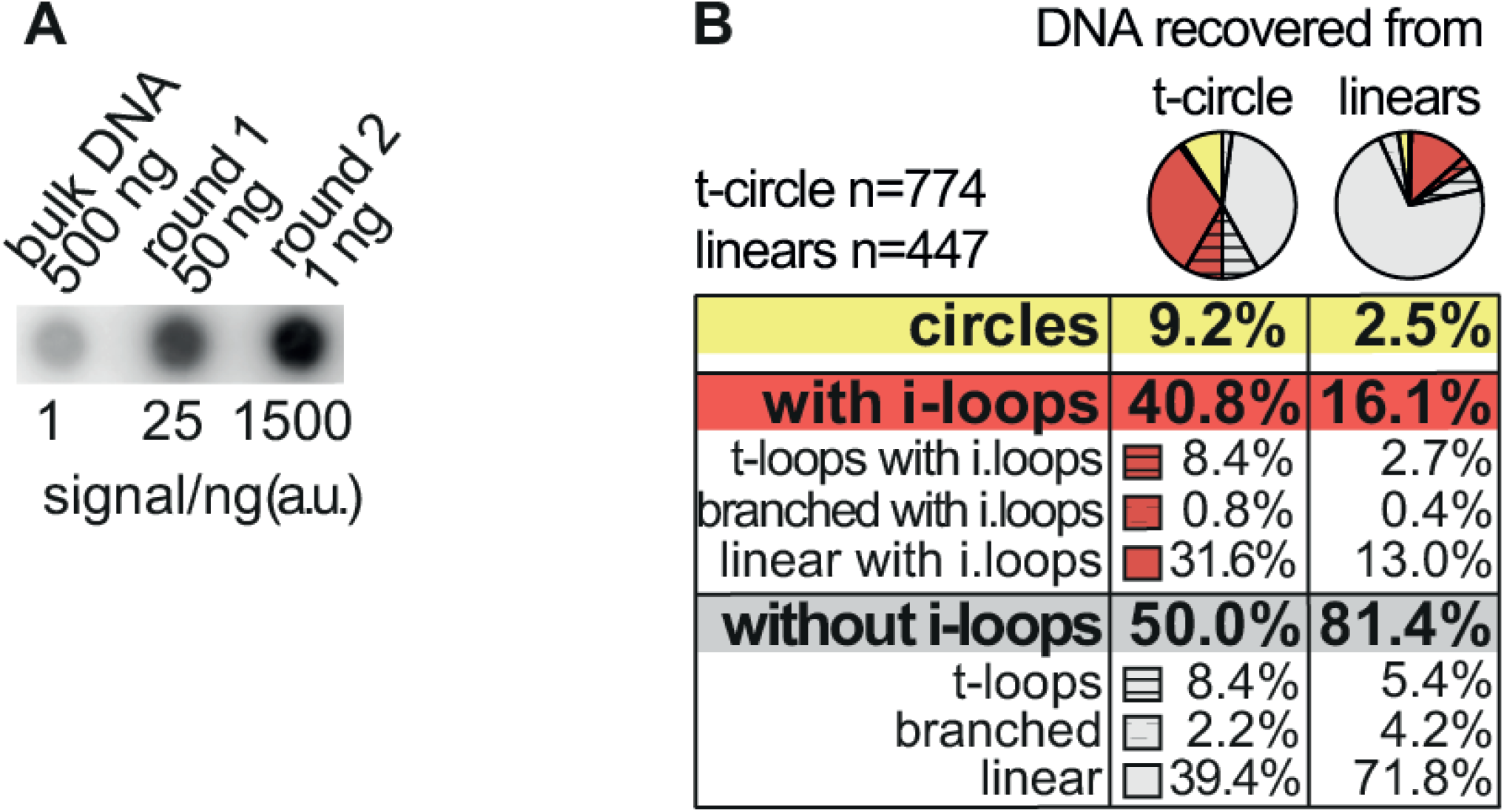
I-loops in the t-circle arc of ALT cells (related to Figure 4). **A.** Dot blot showing the U2OS telomere enrichment for the experiment shown in Figure 4A. The indicated amounts from each enrichment step were spotted on a membrane and hybridized with a probe recognizing the TTAGGG repeats. In the second round, 1 ng of DNA recovered from the linear signal, was used to verify the enrichment. The amount of signal/ng is reported relative to the non-enriched DNA. **B.** This is the same pie chart shown in Figure 4B, that includes the detailed distribution of the molecules recovered from the 2D-gel. Note that i-loops occur also in molecules having a t-loop at the end, or at branched molecules.

**Figure S5.**
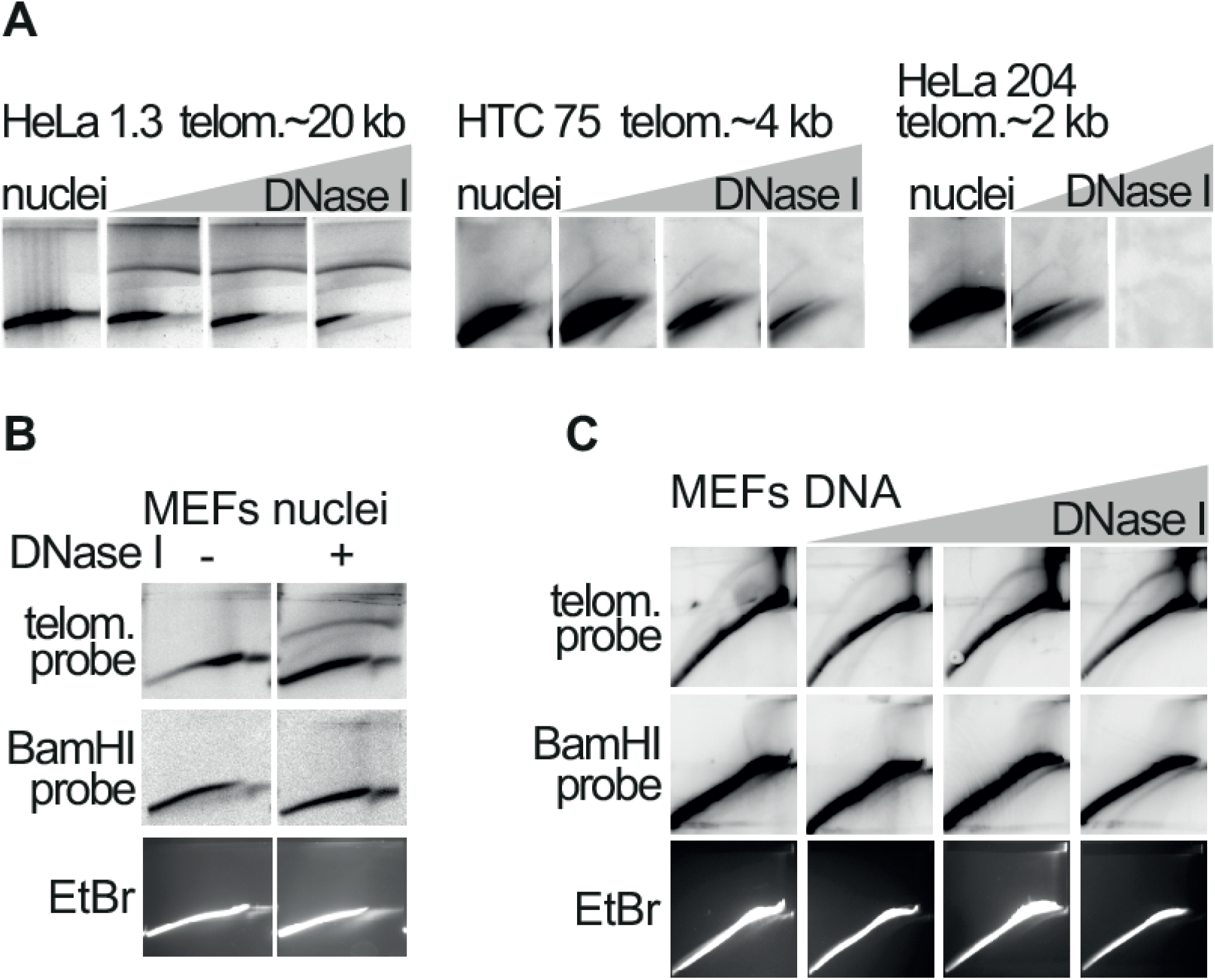
I-loops are induced by single-strand damage at telomeric repeats (related to Figure 5). **A.** 2D-gel analysis showing that the t-circle arc is strongly induced by formation of nicks and gaps at human telomeres. Nuclei prepared from HeLa 1.3 cells (this HeLa clone has long telomeres around, ~20 kb in average) were incubated with either 0; 0.5; 1 or 2.5 μg/ml of DNase I for 8 minutes at RT. Nuclei prepared from HTC75 cells (with telomeres around 4 kb in average) and from HeLa 204 cells (with telomeres around, ~2 kb in average) were incubated with either 0; 5; 10 or 20 μg/ml of DNase I for 8 minutes at RT. The nuclei were then processed for 2D-gel analysis, as described in Figure 5A. **B.** 2D-gel analysis showing that DNase I treatment does not induce the t-circle arc in the bulk DNA or at the BamHI repeats. In a similar experiment as the one described in Figure 5A, nuclei were treated with 2.5 μg/ml of DNase I. Genomic was digested with BglI, split in two and separated on 2D-gels, in duplicate. After blotting, one membrane was hybridized with a probe recognizing the TTAGGG repeats, while the other with a probe recognizing the mouse BamHI repeats. The ethidium bromide staining of one of the second-dimension gels is shown at the bottom. **C.** 2D-gel analysis showing that DNase I treatment on isolated DNA does not induce the t-circle arc in the bulk DNA or at the BamHI repeats. DNA from the same experiment shown in Figure 5B, was digested with KpnI, split in two and then separated in 2D-gels, in duplicate. After blotting, one membrane was hybridized with a probe recognizing the TTAGGG repeats and the other with a probe recognizing the mouse BamHI repeats. The ethidium bromide staining of one of the second-dimension gels is shown at the bottom.

## References

1. Maciejowski, J. & de Lange, T. Telomeres in cancer: tumour suppression and genome instability. Nat Rev Mol Cell Biol 18, 175–186 (2017).

2. Wang, R. C., Smogorzewska, A. & de Lange, T. Homologous Recombination Generates T-Loop-Sized Deletions at Human Telomeres. Cell 119, 355–368 (2004).

3. Pickett, H. A., Cesare, A. J., Johnston, R. L., Neumann, A. A. & Reddel, R. R. Control of telomere length by a trimming mechanism that involves generation of t-circles. EMBO J 28, 799–809 (2009).

4. Tomaska, L., Nosek, J., Kramara, J. & Griffith, J. D. Telomeric circles: universal players in telomere maintenance? Nat Struct Mol Biol 16, 1010–1015 (2009).

5. Henson, J. D. et al. DNA C-circles are specific and quantifiable markers of alternative-lengthening-of-telomeres activity. Nat Biotechnol 27, 1181–1185 (2009).

6. Schmutz, I., Timashev, L., Xie, W., Patel, D. J. & de Lange, T. TRF2 binds branched DNA to safeguard telomere integrity. Nat Struct Mol Biol 24, 734–742 (2017).

7. Tomaska, L., Nosek, J., Kar, A., Willcox, S. & Griffith, J. D. A New View of the T-Loop Junction: Implications for Self-Primed Telomere Extension, Expansion of Disease-Related Nucleotide Repeat Blocks, and Telomere Evolution. Front Genet 10, 792 (2019).

8. Zellinger, B., Akimcheva, S., Puizina, J., Schirato, M. & Riha, K. Ku suppresses formation of telomeric circles and alternative telomere lengthening in Arabidopsis. Mol Cell 27, 163–169 (2007).

9. Deng, Z., Dheekollu, J., Broccoli, D., Dutta, A. & Lieberman, P. M. The origin recognition complex localizes to telomere repeats and prevents telomere-circle formation. Curr Biol 17, 1989–1995 (2007).

10. Wang, Y., Ghosh, G. & Hendrickson, E. A. Ku86 represses lethal telomere deletion events in human somatic cells. Proc Natl Acad Sci U S A 106, 12430–12435 (2009).

11. Gu, P. et al. CTC1 deletion results in defective telomere replication, leading to catastrophic telomere loss and stem cell exhaustion. The EMBO Journal 31, 2309–2321 (2012).

12. O’Sullivan, R. J. et al. Rapid induction of alternative lengthening of telomeres by depletion of the histone chaperone ASF1. Nat Struct Mol Biol 21, 167–174 (2014).

13. Li, J. S. et al. TZAP: A telomere-associated protein involved in telomere length control. Science 355, 638–641 (2017).

14. Zhang, T. et al. Strand break-induced replication fork collapse leads to C-circles, C-overhangs and telomeric recombination. PLoS Genet 15, e1007925 (2019).

15. Doksani, Y. The Response to DNA Damage at Telomeric Repeats and Its Consequences for Telomere Function. Genes (Basel) 10, (2019).

16. Cohen, S. & Lavi, S. Induction of circles of heterogeneous sizes in carcinogen-treated cells: two-dimensional gel analysis of circular DNA molecules. Mol Cell Biol 16, 2002–2014 (1996).

17. Cesare, A. J. & Griffith, J. D. Telomeric DNA in ALT Cells Is Characterized by Free Telomeric Circles and Heterogeneous t-Loops. Molecular and Cellular Biology 24, 9948–9957 (2004).

18. de Lange, T. et al. Structure and variability of human chromosome ends. Mol Cell Biol 10, 518–527 (1990).

19. Griffith, J. D. et al. Mammalian telomeres end in a large duplex loop. Cell 97, 503–514 (1999).

20. Lopes, M. Electron microscopy methods for studying in vivo DNA replication intermediates. Methods Mol Biol 521, 605–631 (2009).

21. Nikitina, T. & Woodcock, C. L. Closed chromatin loops at the ends of chromosomes. Journal of Cell Biology 166, 161–165 (2004).

22. Nabetani, A. & Ishikawa, F. Unusual telomeric DNAs in human telomerase-negative immortalized cells. Mol Cell Biol 29, 703–713 (2009).

23. Riley, D. E. Deoxyribonuclease I generates single-stranded gaps in chromatin deoxyribonucleic acid. Biochemistry 19, 2977–2992 (1980).

24. Hanson, C. V., Shen, C. K. & Hearst, J. E. Cross-linking of DNA in situ as a probe for chromatin structure. Science 193, 62–64 (1976).

25. Sfeir, A. et al. Mammalian Telomeres Resemble Fragile Sites and Require TRF1 for Efficient Replication. Cell 138, 90–103 (2009).

26. Crabbe, L., Verdun, R. E., Haggblom, C. I. & Karlseder, J. Defective telomere lagging strand synthesis in cells lacking WRN helicase activity. Science 306, 1951–1953 (2004).

27. Du, X. et al. Telomere shortening exposes functions for the mouse Werner and Bloom syndrome genes. Mol Cell Biol 24, 8437–8446 (2004).

28. Vannier, J. B., Pavicic-Kaltenbrunner, V., Petalcorin, M. I., Ding, H. & Boulton, S. J. RTEL1 Dismantles T Loops and Counteracts Telomeric G4-DNA to Maintain Telomere Integrity. Cell 149, 795–806 (2012).

29. Gaubatz, J. W. & Flores, S. C. Tissue-specific and age-related variations in repetitive sequences of mouse extrachromosomal circular DNAs. Mutat Res 237, 29–36 (1990).

30. Sinclair, D. A. & Guarente, L. Extrachromosomal rDNA circles--a cause of aging in yeast. Cell 91, 1033–1042 (1997).

31. Rivera, T., Haggblom, C., Cosconati, S. & Karlseder, J. A balance between elongation and trimming regulates telomere stability in stem cells. Nat Struct Mol Biol 24, 30–39 (2017).

32. Vollenweider, H. J., Sogo, J. M. & Koller, T. A routine method for protein-free spreading of double- and single-stranded nucleic acid molecules. Proc Natl Acad Sci U S A 72, 83–87 (1975).

33. Pipkin, M. E. & Lichtenheld, M. G. A reliable method to display authentic DNase I hypersensitive sites at long-ranges in single-copy genes from large genomes. Nucleic Acids Res 34, e34 (2006).

34. de Lange, T. Human telomeres are attached to the nuclear matrix. EMBO J 11, 717–724 (1992).

35. Fanning, T. G. Size and structure of the highly repetitive BAM HI element in mice. Nucleic Acids Res 11, 5073–5091 (1983).

